# A Multi-Omics Study of Cognitive Resilience to Alzheimer’s Disease

**DOI:** 10.64898/2026.09.02.746746

**Authors:** Erming Wang, Boer Xie, Minghui Wang, Yingxue Fu, Xusheng Wang, Yuxin Li, Peng Xu, Huaping Liu, Lap Ho, Zhidong Tu, Charles Mobbs, Chris Gaiteri, Michelle E. Ehrlich, David A. Bennett, Vahram Haroutunian, Junmin Peng, Bin Zhang

## Abstract

Alzheimer’s disease (AD) is the most severe form of dementia. While significant efforts have been made to identify AD risk factors and develop therapeutics, much less attention has been given to neurobiological factors and molecular signatures associated with cognitive resilience. Substantial evidence suggests that modifiable factors, such as physical and mental activity, may contribute to cognitive reserve and resilience. To this end, we developed a resilient cohort of paired genetics, transcriptomics and proteomics data from the prefrontal cortex of 282 cognitively normal older adults. We conducted extensive preprocessing and rigorous quality control (QC), including QC on sequence metrics and sample matching across multiomics. This cohort can be leveraged for identification of molecular signatures of cognitive resilience to AD and is freely shared with the research community.

## Introduction

Ageing is the greatest known risk factor for the development of Alzheimer’s disease (AD)^1^. Up to 50% of persons of 85 years or older are clinically demented and the estimated healthcare costs continue to increase along with the aging population^2^. However, a large proportion of old individuals show successful aging, i.e., without significant signs of cognitive decline^3^. While considerable effort has been expanded to identify AD risk factors and developing treatment, less attention has been given to neurobiological factors and molecular signatures associated with cognitive resilience. Identification of neurobiological factors and genetic networks associated with cognitive resilience could reveal potential druggable targets towards prevention, rather than merely treatment or reversal, of cognitive decline.

Substantial evidence suggests that modifiable factors, such as education level, occupation, and physical and mental activity, may contribute to cognitive reserve and resilience ^4–8^. However, the molecular mechanisms underlying cognitive resilience remain elusive. Markers of cognitive reserve and resilience in aged individuals have mainly based on imaging data, including morphometry, volume, connectivity, and ventricular size^9,10^ as well as cerebrospinal Aβ-42 and Tau levels^11^. Attempts were made to investigate the associations between imaging markers and genetic factors related to longevity and resilience. For example, the longevity gene *KLOTHO* has been associated with increased volume of the dorsolateral prefrontal cortex (DLPFC), a region important for executive function^12^.

Molecular and genetic factors contributing to resilience in elders show complex interactions and overlap with those associated with longevity and they are involved in pathways that may influence the same molecular networks implicated in aging and/or AD, but in an opposite manner^13,14^. The known resilience-related pathways include synaptic integrity and plasticity^15^, immune competence^16^, bioenergetics^17,18^, cell death and cell-cycle regulation^19,20^, angiogenesis and heme biosynthesis^20^. Multiomics analysis of brain samples from the population resilient to AD will enable unbiased identification of molecular factors underlying resilience. To this end, we have developed a resilient cohort of matched genetics, transcriptomics and proteomics data from the prefrontal cortex of 282 cognitively normal adults (64∼101 years) selected from the Mount Sinai/JJ Peters VA Medical Center Brain Bank^21,22^ (MSBB) and the Religious Orders Study/Memory and Aging Project^23–27^ (ROSMAP). We have conducted extensive preprocessing and rigorous quality control (QC), including QC on sequence metrics and sample matching across multiomics. This cohort, which is shared with the research community, can be leveraged for identification of molecular signatures of cognitive resilience to AD.

## Methods

### Human tissues

We collected frozen postmortem tissue excised from the PFC region of 282 normal control brains from the MSBB^21,22^ (n = 82) and the ROSMAP^7,23,28,29^ (n = 200) cohorts (**Figure 1A**). The neuropsychological, diagnostic and autopsy protocols of the MSBB participants were reviewed and approved by Institutional Review Boards of the Mount Sinai and JJ Peters VA Medical Center (IRB# 13-00709 and HAR-13-059). Informed consent was received from all MSBB participants or their representatives. ROSMAP studies were approved by an Institutional Review Board of Rush University Medical Center (ROS IRB# L91020181, MAP IRB# L86121802). All ROSMAP participants were enrolled without known dementia and agreed to detailed clinical evaluation and brain donation at death. The ROSMAP studies were conducted according to the principles expressed in the Declaration of Helsinki. Each participant signed an informed consent, Anatomic Gift Act, and an RADC Repository consent (IRB# L99032481) allowing the data and biospecimens to be repurposed.

**Figure 1.**
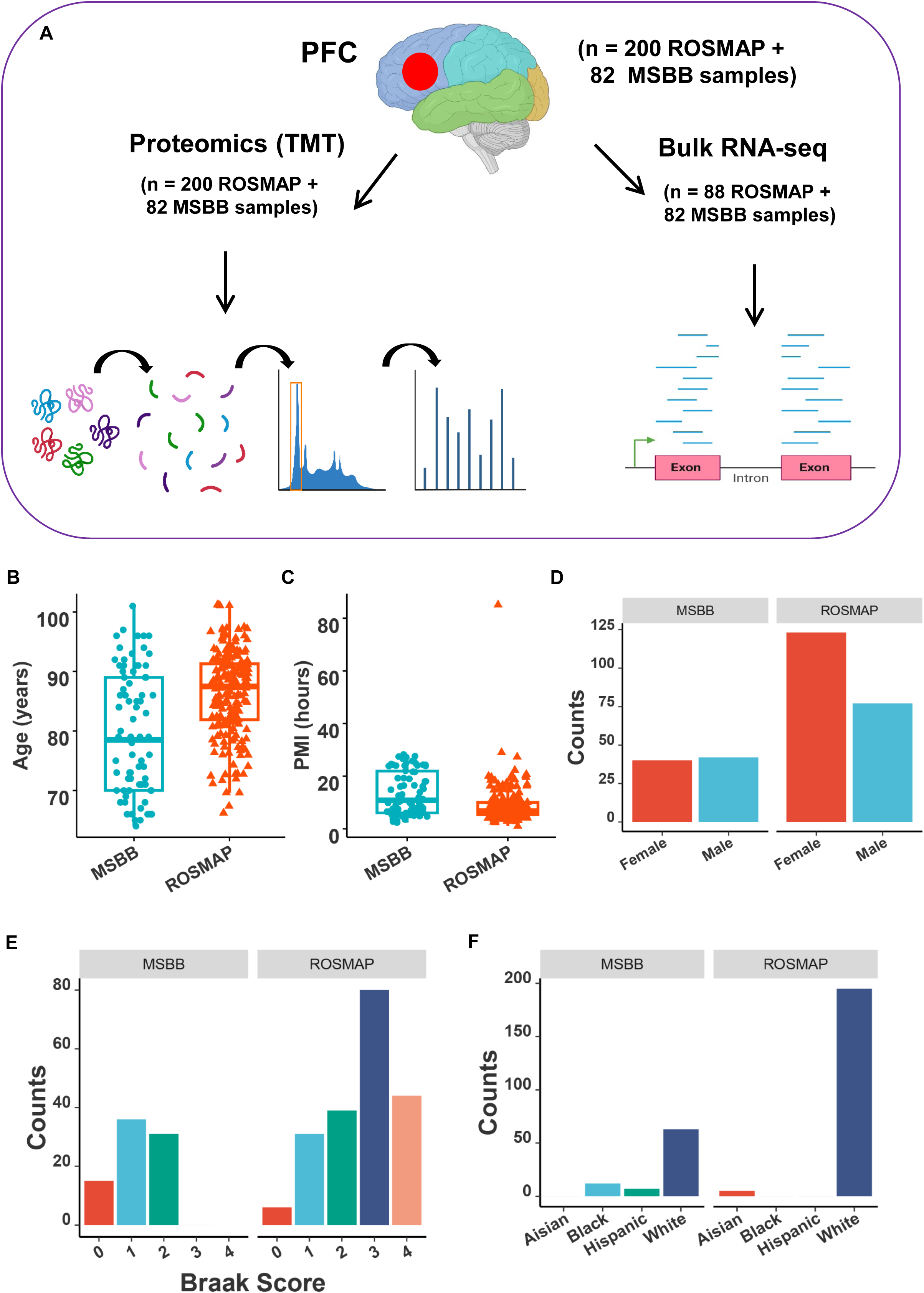
Schematic representation of study design, and summary of demographic and clinical traits of human samples. A, study design. Human brain postmortem tissue was dissected from the prefrontal cortex (PFC) region, and subject to bulk tandem mass tag- (TMT-) proteomics profiling (left panel) and bulk RNA-seq assays (right panel). B, boxplots to show the distribution of age at death (AOD) across each cohorts. C, boxplots to show the distribution of postmortem interval (PMI) across each cohorts. D, barplots to show the numbers of subjects over female vs male across each cohorts. E, barplots to show the numbers of subjects over different Braak score level across each cohorts. F, barplots to show the race distribution score level across each cohort. Some of the diagrams were adapted from BioRender.

The subjects had an average age at death of 79 (ranging from 64 to 101) and 86 (ranging from 66 to 101) years for the MSBB and ROSMAP samples, respectively (**Figure 1B**). Most subjects (93%) had a postmortem interval (PMI) less than 24 hours (**Figure 1C**). The subjects had a ratio of 40:42 and 123:77 in females vs males for the MSBB and ROSMAP samples, respectively (**Figure 1D**). All of the subjects had a low or medium severity in Alzheimer’s disease (AD) tau pathology as determined by Braak scores^26,30^ (**Figure 1E**). In addition, most of the subjects are white^31,32^ (**Figure 1F**). Detailed demographic and neuropathological assessments as well as cognitive and functional data are available for all donors (see Data Records).

All 282 samples were subject to proteomics profiling, which is only reported in this work. Among the 282 samples, 112 had RNA-seq from the previous sequencing effort by the Rush Alzheimer’s disease center (RADC) (https://www.synapse.org/Synapse:syn3388564) (**Figure 2A**). Therefore, bulk RNA-seq data were generated for the remaining 170 samples in this study, including all the 82 samples from the Mount Sinai Brain Bank (MSBB) and 88 samples from the ROSMAP cohort (**Figure 2A**). Out of the 282 samples, 226 (80.1%) went through whole genome sequencing (WGS) previously, including 157 ROSMAP samples from the AMP-AD project (https://www.synapse.org/Synapse:syn10901595) and 69 MSBB samples from the National Institute of Mental Health Data Archive (NDA) repository (collection 3917) (**Figure 2A**).

**Figure 2.**
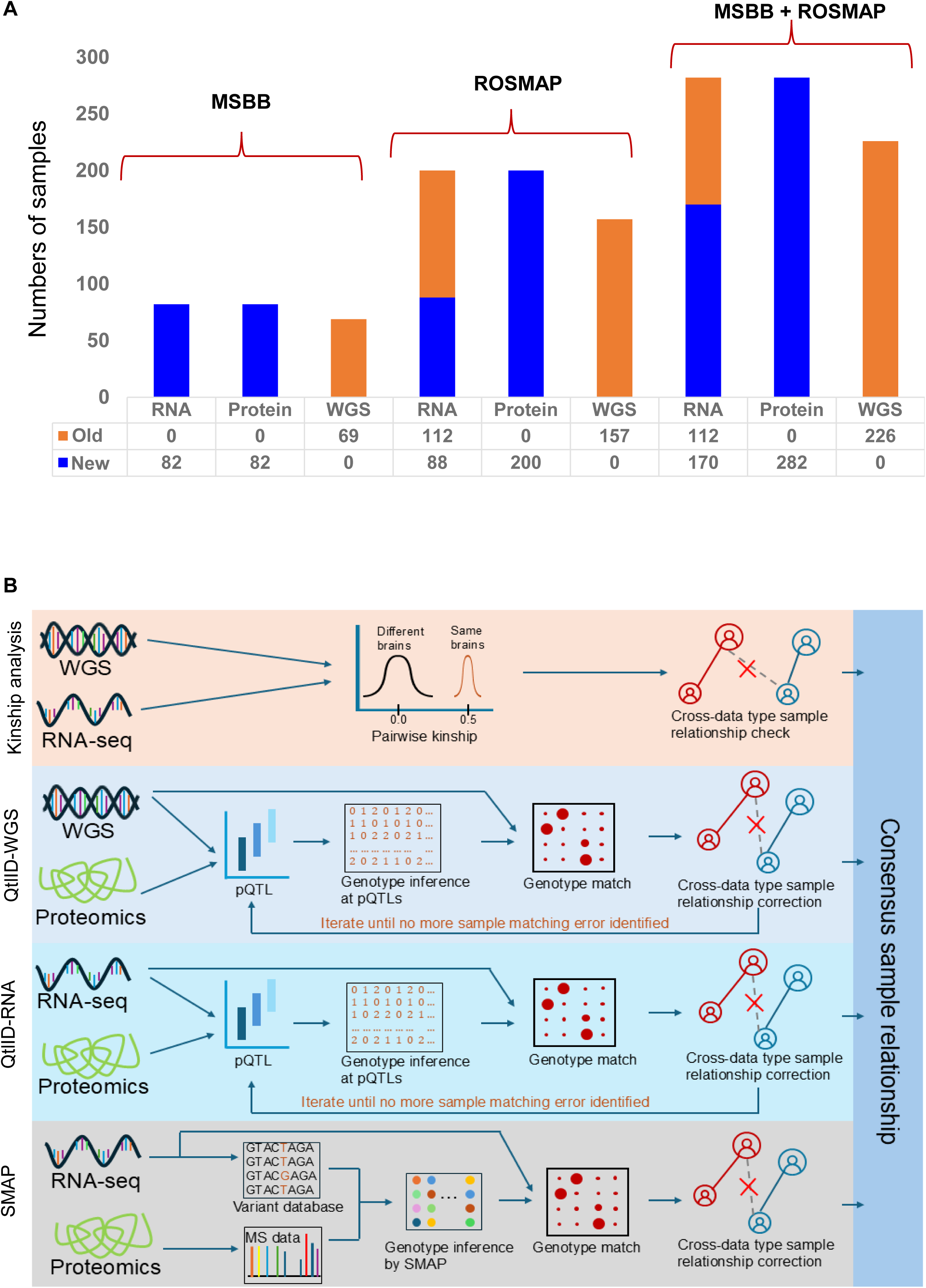
Summary of samples in multi-omics and outline of quality control workflow. A, summary of samples across various omics profiling. New and old stand for sequencing/profiling conducted in this study and previous studies, respectively. RNA, Protein, and WGS stand for bulk RNA-seq, proteomics profiling, and whole genome sequencing, respectively. B, the robust and efficient pipeline for quality control and matching multiple-omics data across modalities from the same cohort. Each color block of the left panel indicates a step in matching one data type to another. From top to bottom are matching between WGS and RNA-seq using kinship analysis, matching between WGS and proteomics using pQTLs (QtlID-WGS), matching between RNA-seq and proteomics by two independent approaches (QtlID-RNA, and sample matching in proteogenomics (SMAP)), respectively. Lastly, the pair-wise data type matchings in the left panel are cross-compared to derive the final consensus sample identity relationship verification (the right panel).

### Bulk RNA-seq

We have performed bulk RNA-seq assays on 170 brains, out of which 82 and 88 brains are from the MSBB and ROSMAP cohorts, respectively. The results of these bulk RNA-seq assays have not been reported elsewhere before.

#### RNA extraction

Total RNA was extracted from flash frozen post-mortem tissue of 30 mg. Trizol/Chloroform extraction method (Trizol catalog #15-596-018, Invitrogen) was used, followed by Qiagen RNeasy minikit column purification. The column purification step was used to ensure the quality of extracted RNA.

#### Library preparation and sequencing

RNA libraries were prepared using the KAPA Stranded RNA-seq with RiboErase (H/M/R) library preparation kit (Roche 07962304001) in accordance with the manufacturer’s instructions. 500ng of total RNA samples were ribodepleted using oligonucleotide hybridization and RNase H treatment followed by DNase treatment. The RNA was fragmented using divalent cations under elevated temperature. The cleaved RNA fragments were copied into cDNA complementary molecules, adenylated, ligated to Illumina sequencing adapters, purified and enriched with PCR to create the final cDNA library. Final libraries were quantified using the Qubit Fluorometer (Life Technologies) or Spectramax M2 (Molecular Devices) and evaluated for size distribution on the Fragment Analyzer (Agilent). Libraries were sequenced on NovaSeq 6000 (Illumina) with 100bp paired-end reads.

#### Computation methods

Sequencing quality control was performed using Picard^33^ (version 1.83) and RSeQC^34^ (version 2.6.1). STAR (version 2.5.2a)^35^ was used to align reads to the GRCh37 genome, using Gencode v25 annotation. The raw RNA count data was further processed to remove entries with low counts, and then normalized and adjusted for covariables following the in-house workflow as described in^36,37^.

### Bulk Tandem mass tag- (TMT-) proteomics profiling

#### Tandem mass tag analysis

Following our optimized 11-plex TMT protocol^38–43^, ∼10 mg of brain tissue from each individual sample was weighed upon arrival, lysed, and analyzed for protein quality. The 282 samples were organized into 29 batches using block randomization to account for confounding factors such as AD-resilient vs. healthy status, gender, and age. An internal standard was generated by pooling a small fraction from all samples and included in every batch. For each batch, proteins were digested, labeled with TMT reagents, and pooled for two-dimensional liquid chromatography-tandem mass spectrometry (LC/LC-MS/MS) analysis. The first dimension of LC was used to reduce sample complexity, improving protein identification, while the second dimension was directly coupled with MS/MS for both peptide/protein identification and quantification.

#### Computation methods

Proteomics data analysis was performed as previously described^22,44,45^. The hybrid JUMP search engine^44^ was used to search TMT MS/MS raw data against a composite target/decoy database^24^ to evaluate false discovery rate (FDR). The protein database was constructed by combining downloaded SwissProt, TrEMBL, and UCSC databases, followed by redundancy removal, resulting in 83,955 entries for human proteins. This target database was concatenated with a decoy database to enable FDR estimation. Key parameters for the search included a mass tolerance of 15 ppm for precursor ions and 10 ppm for product ions, full tryptic specificity, static modifications of TMT tags (+229.162932 Da) on lysine residues and peptide N-termini, as well as carbamidomethylation on cysteine (+57.02146 Da). Dynamic modifications included methionine oxidation (+15.99492 Da), with a maximum of two missed cleavage sites and up to three modification sites allowed. Peptide-spectrum matches (PSMs) were filtered based on precursor ion mass accuracy and minimal search score. PSMs were then grouped by peptide length, tryptic ends, modifications, missed cleavages, and precursor ion charge states. Further filtering using JUMP-based matching scores (Jscore and ΔJn) was applied to achieve 1% protein FDR. In cases where a peptide could be assigned to multiple homologous proteins, the rule of parsimony was applied, and the peptide was assigned to the canonical form of the protein in the SwissProt database. If no canonical form was defined, the peptide was assigned to the protein with the highest number of PSMs.

Proteins were quantified in the following steps, similar to previous reports^22,25,26^: (i) the signals of TMT reporter ions were identified in each PSM; (ii) the signals were corrected according to isotopic distribution of TMT reagents; (iii) PSMs with extremely weak signals were discarded (e.g. minimum signal < 1,000 and median signal < 5,000 in one PSM); (iv) sample loading bias was normalized by the trimmed median signal of all PSMs; (v) for protein relative quantification, the PSM signal ratios were summarized into protein signal ratios; (vi) for protein absolute quantification, the relative signal ratios were multiplied by the grand-mean intensity of the top three most highly abundant PSMs from the protein. In addition, we performed y_1_-ion based correction of TMT data^25^. To combine all quantification tables for different batches, the internal standard was used for batch normalization. The raw data and processed data were deposited in Synapse^27^.

### Cross-data sample matching

The present resilience study cohort includes matched RNA-seq and proteomics data from each subject, as well as WGS data for a majority (226 out of 282 samples) of the subjects (**Figure 2A**). Given the scale and complexity of the dataset, sample errors, including incorrect labeling, sample swapping and contamination, are inevitable but difficult to detect by the platform specific QC procedure on each type of data. Systems approach for QC across multiple-omics data types is essential for reliable technical validation such that true biological signals can be accurately uncovered from the integrative data analysis. To this end, we applied a well-established computational pipeline to identify the consistent sample mapping across the multiple types of molecular data^46,47^. As illustrated in **Figure 2B** and detailed below, this pipeline first matches WGS and RNA-seq samples using genotype-based kinship analysis, and then matches proteomics samples with either WGS or RNA-seq samples through protein quantitative trait loci (pQTLs)-based genotype inference approach^21,48^. For simplicity, we called the two pQTL-based proteomics sample matching approaches by QtlID-WGS and QtlID-RNA for matching WGS or RNA-seq samples, respectively. In parallel, a complementary proteogenomic approach (sample matching in proteogenomics (SMAP^45^)) was employed to infer sample-specific peptide-level mutations based on RNA-seq variants database for matching proteomics samples with RNA-seq samples. Lastly, a consensus proteomics sample identity calling was derived by comparing the 3 sets of proteomics sample identity inferences (**Figure 2B**).

### Gene set enrichment analysis

The Peason’s correlation between age at death (AOD) and individual genes or proteins was first calculated. The gene list was determined to have correlation with a nominal p < 0.05, and then ranked by the correlation coefficients in descending order. The protein sets were obtained by splitting the AOD-associated protein signature (p.adj < 0.05) into negatively- and positively-associated protein signatures. The gene list was then queried for enrichment over the negatively- and positively-associated protein signatures, respectively using the function GSEA in the R package clusterProfiler^49^.

### Data Records

All the datasets presented herein have been deposited onto the AMP-AD Knowledge Portal with a specific synapse id of syn52676457 (https://www.synapse.org/Synapse:syn52676457). These include raw sequencing data for bulk RNA-seq and raw proteomics profiling data along with raw sample metadata. We also provided processed RNA-seq and proteomics expression data after quality control (QC), as well as aligned corrected sample metadata.

### Technical assessment Bulk RNA-seq

Bulk RNA-seq assays were conducted on 82 and 88 tissue samples for the MSBB and ROSMAP cohorts, respectively. We assessed the quality of the RNA-seq data using the following metrics: RNA integrity number^50^ (RIN) (**Figure 3A**), ribosomal RNA rate (**Figure 3B**), GC content (**Figure 3C**), total number of reads (**Figure 3D**), mapping rate (**Figure 3E**), uniquely mapping rate (**Figure 3F**), coding sequence (CDS) rate (**Figure 3G**), exonic rate (**Figure 3H**), and intergenic rate (**Figure 3I**). Most of the samples (78.8%) possessed an acceptable RIN value (> 4.0), whereas only 5 samples had a RIN less than 2 (**Figure 3A**).

**Figure 3.**
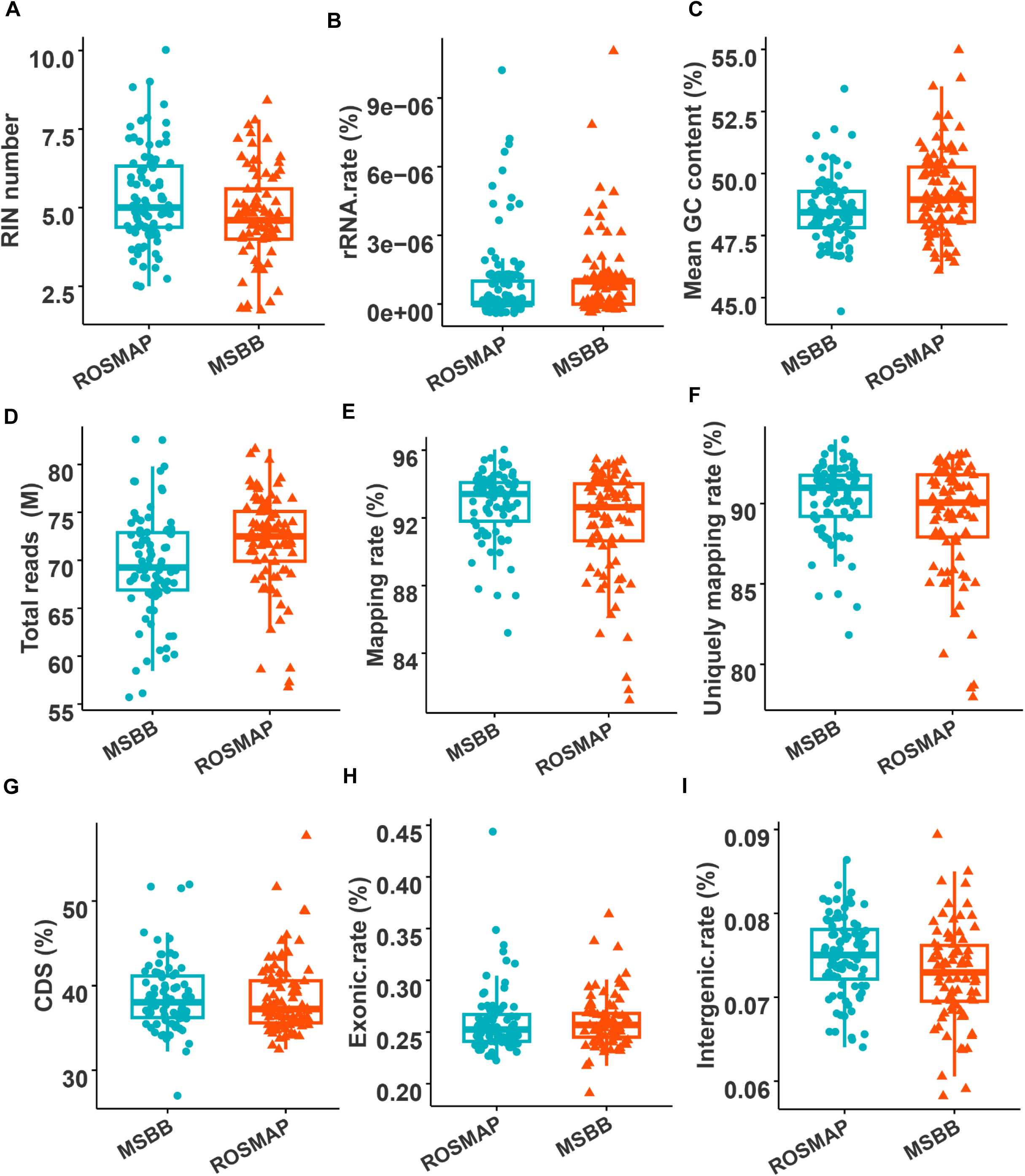
Summary of demographic and clinical traits, and analysis of assay metrics for the RNA-seq samples. A-I, barplots to show RNA-seq assay QC metrics: RIN (A), ribosomal RNA rate (B), average GC content (C), total number of reads (D), mapping rate (E), uniquely mapping rate (F), CDS rate (G), exonic rate (H), and intergenic rate (I).

We estimated the proportion of variation in gene expression^51^ that could be explained by various biological and technical covariates (**Figure 4A**). As shown in **Figure 4A**, the RIN is the most prominent covariate, followed by exonic rate, intergenic rate, project (MSBB vs ROSMAP), age at death and RNA rate, in contrast to the inferior contribution from PMI, Braak score, race and sex. We then removed the effects from the covariates (RIN, exonic rate, intergenic rate, project (MSBB vs ROSMAP), ribosomal RNA rate, PMI, and race) from the gene expression essentially following the method as described in^36,37^. Depending on the downstream analysis, age at death and/or sex would also be treated as a covariate.

**Figure 4.**
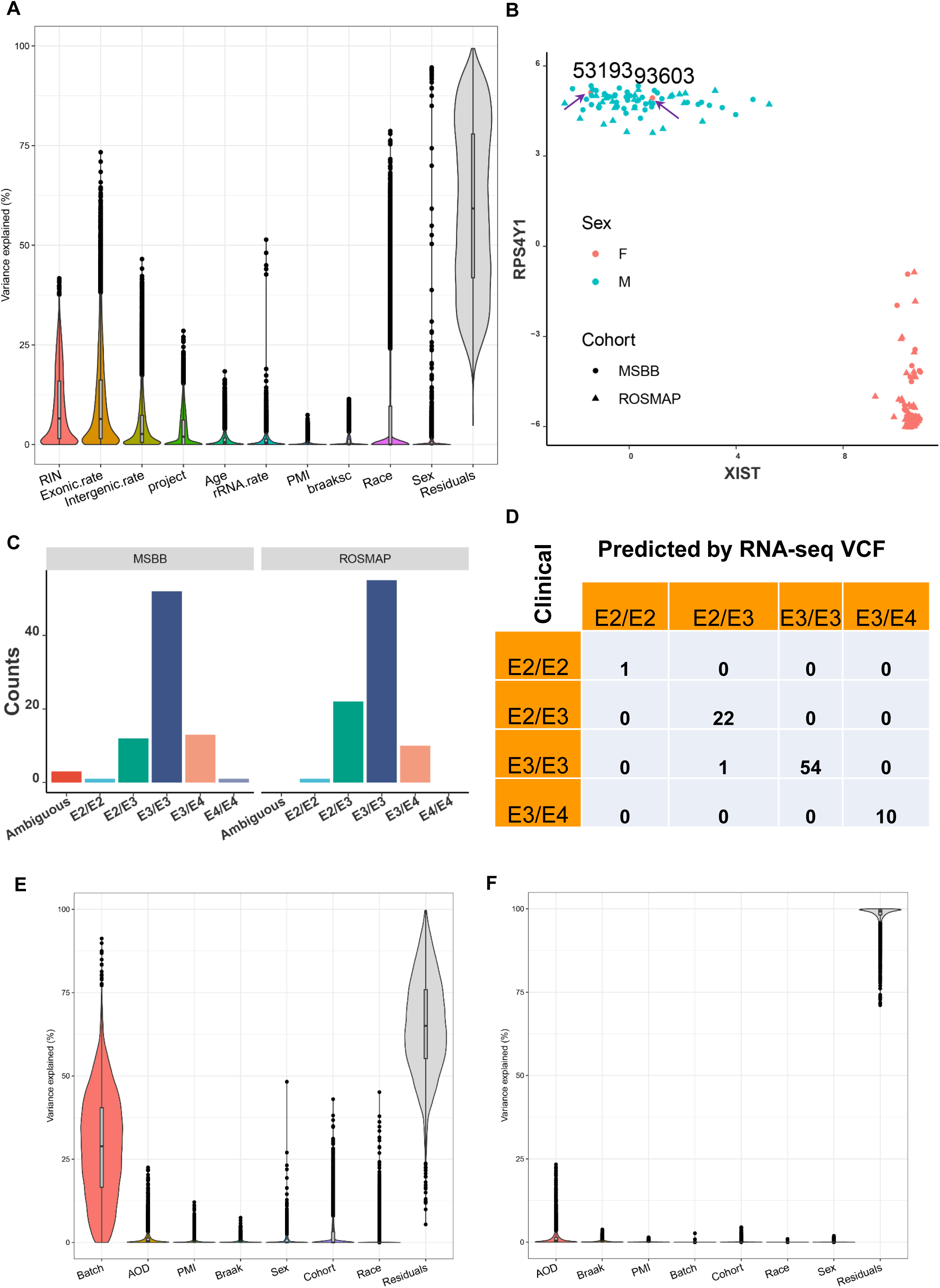
Analysis of assay variance and genetic similarity. A, variance explained by biological and technical covariates for bulk RNA-seq expression. B, sex verification via quantification of the expression of male-(RPS4Y1) and female-specific genes (XIST). Arrows point to the samples potentially mislabeled. Text labels above show the subject IDs for the two potentially mislabeled samples. C-D, APOE genotype prediction (C) and validation (D). E-F, variance explained by biological and technical covariates for proteomics expression before (E) and after (F) the covariable adjustment. Note, in F, the variable AOD was not included in the covariable adjustment. Note, A-D are the results from bulk RNA-seq assays, whereas E-F are for proteomics profiling.

To verify the expression of genes on sex chromosomes is consistent with the reported sex, we monitored the expression of the female sex-specific XIST in contrast to that of the male-specific gene RPS4Y1^21,22^ (**Figure 4B**). As shown in **Figure 4B**, all the samples are clustered according to their reported sex labels except for two samples in the male cluster, implying potential mislabeling of these two samples from male to female sex. We also inferred the APOE genotype for each of the samples using the per-sample variants determined from raw RNA-seq sequencing^52^ (**Figure 4C-D**). Since we have the laboratory-reported APOE genotypes for the ROSMAP samples, we compared their concordance between the RNA-seq-derived and laboratory-reported APOE genotypes. We found that 87 out of the 88 ROSMAP samples had a consistent identity between the RNA-seq-derived and laboratory-reported APOE genotype (**Figure 4D**).

Together, the above-QC results indicated that an overall good quality was achieved in the RNA-seq assay. The detected identity swaps between samples and sex-mislabeling were corrected, and duplicated samples were removed by leaving only one sample in the processed expression data and sample meta data.

### Proteomics profiling

We identified 11,803 protein isoforms (thereafter, termed proteins) that have detectable expression in more than 50% of the 282 samples, which spans over 9,462 genes. Missing values were imputed using the method as described in^53^, and protein expression normalization was performed by log2-transformation.

We then estimated the proportion of variation in protein abundance^51^ that could be explained by various biological and technical covariates (**Figure 4E**). As shown in **Figure 4E**, batch is the most prominent covariate, followed by age at death, in contrast to the inferior contribution from PMI, Braak score, sex, cohort (MSBB vs ROSMAP) and race. We further removed the effects from the covariates (batch, cohort (MSBB vs ROSMAP), PMI, and race) from the protein expression (**Figure 4F**) using a mixed linear model following the method as described in^37^. Depending on the downstream analysis, age at death (AOD) would also be treated as a covariate and corrected from the protein expression.

### Sample matching between WGS and RNA-seq

To match WGS and RNA-seq data, we leveraged existing WGS-based genotyping data for MSBB samples deposited in the National Institute of Mental Health Data Archive (NDA) and for ROSMAP samples deposited in the AMP-AD knowledge portal. For resilience RNA-seq samples, we called genetic variants using established in-house pipeline following the genome analysis toolkit (GATK) best practices^46^. We estimated pairwise sample kinship using KING^54^ to compare the genetic concordance among all sequencing samples between WGS and RNA-seq data (**Figure 5A**). Based on the kinship analysis, 3 RNA-seq samples were found to be mislabeled, and another 6 RNA-seq samples failed to match any WGS sample at 2nd degree relationship. We removed the 6 RNA-seq samples that failed to match WGS data from matching with proteomics data (**Supplementary Table 1**).

**Figure 5.**
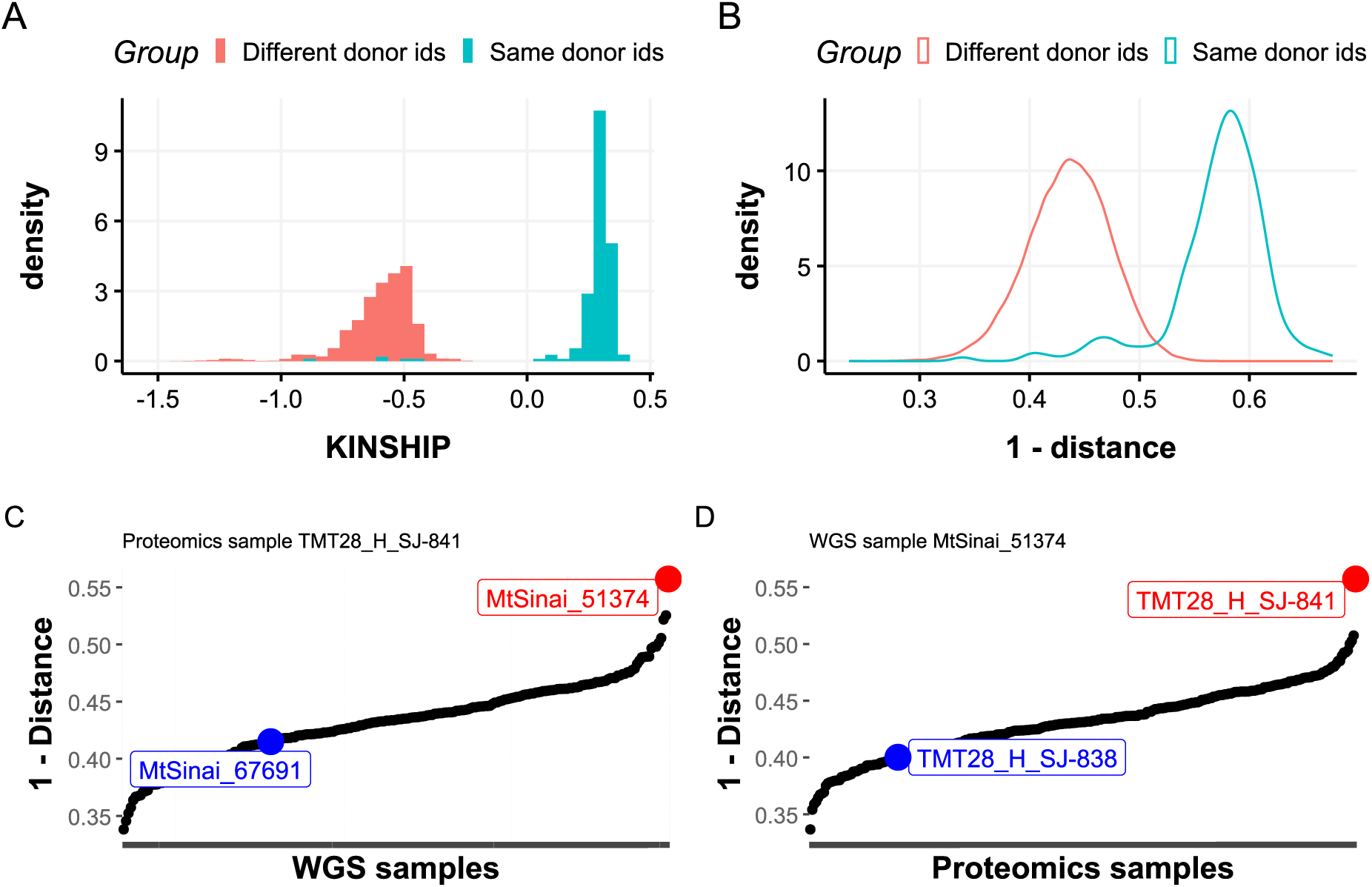
Identification of mis-labeled samples via genetic inference and similarity test across data types. A, Distribution of pairwise sample kinship estimates between WGS and RNA-seq. B. Distribution of pairwise sample genetic similarities between WGS and proteomics samples. C-D, Genetic similarity check reveals a pair of potentially mislabeled proteomics and WGS samples. C, Sorted sample genetic similarity across WGS samples for proteomics sample TMT28_H_SJ-8416. D, Sorted sample genetic similarity across proteomics samples for WGS sample MtSinai_51374. Green and red colors highlight the expected match by annotation table and the observed best match inferred by the QtlID-WGS approach.

### pQTL-based sample matching between proteomics and WGS/RNA-seq

To match WGS and proteomics samples, our QtlID-WGS approach (**Figure 2B**) first computed protein QTLs (pQTLs) based on protein expression and WGS genotype data and then inferring genotypes at the cis-SNPs to compare against the observed WGS genotypes at these loci, such that we could compute the genetic similarities of all possible pairs between WGS and proteomics samples as we previously did^46,47^. **Figure 5B** indicates a clear separation of proteomics-WGS pairwise sample similarity measures between pairs annotated with the same donor and pairs annotated with different donors. A pair of proteomic and WGS samples was considered to be properly matched if 1) they were mutually best matched with each other across all samples, and 2) they had the same donor id by annotation. On the other hand, a pair of proteomic and WGS samples was considered to be potentially mislabeled if 1) they were mutually best matched with each other across all samples, and 2) they had different donor ids by annotation. **Figures 5C-D** show an example of a potentially mislabeled pair of proteomics and WGS samples. While proteomics sample TMT28_H_SJ-8416 was expected to match WGS sample MtSinai_67691 by annotation, its mutual best matched WGS sample was MtSinai_51374. When potential mislabeling was identified, we considered the error occurred in the proteomics samples since we observed similar mislabelings when matching proteomics with RNA-seq. Then we corrected the sample label(s) and repeated the sample matching between the WGS data and newly updated proteomics data. This sample match and correction procedure was repeated until no improvement in sample matching could be made.

Analogously, we leveraged RNA-seq genotype data to conduct pQTL-based sample matching between proteomics and RNA-seq data for donors with both types of data (QtlID-RNA, **Figure 2B**). No disagreement in proteomics sample identity mappings was identified for proteomics samples that can be successfully mapped by both the QtlID-WGS and the QtlID-RNA approaches.

### SMAP-based sampling matching between proteomics and RNA-seq

Additionally, we utilized a proteogenomics approach to identify sample-specific peptides with mutations, followed by the proteomics genotype inference and sample alignment with RNA-seq using the SMAP software^47^. Briefly, by constructing customized protein database using SNVs detected from RNA-seq data, peptides with sample specific mutations were identified using the JUMPg software^48^. The resulting peptides with mutations were quantified and processed by SMAP for sample alignment with two steps: (i) inference of sample specific genotype based on TMT-based quantification while taking the genotype dosage information in the RNA-seq data as prior knowledge; and (ii) sample verification and correction by comparing the inferred genotypes versus the mutation profiles of the RNA-seq samples with high quality genotypic data.

### Consensus calling of proteomics sample identity verification

To derive the final verification of proteomics sample identities, we compared the proteomics sample identity (ID) inferences across three approaches (i.e. QtlID-WGS, QtlID-RNA, and SMAP). As WGS variants were more accurate than RNA-seq variants, we considered QtlID-WGS approach to be the most accurate when inconsistency occurred. However, it is noted that not all proteomics samples had WGS data. As a result, among the 281 unique proteomics samples (2 duplicates from the same donor), 266 had donor identities successfully verified. These included 186 proteomics samples consistently inferred by all three approaches, 2 proteomics samples consistently inferred by QtlID-WGS and QtlID-RNA, 10 proteomics samples consistently inferred by QtlID-WGS and SMAP, 4 proteomics samples inferred by QtlID-WGS only, and 64 consistently inferred by both QtlID-RNA & SMAP. The remaining 15 proteomics samples had their donor identities undetermined due to the lack of high quality WGS or RNA-seq variants data (**Supplementary Table 2**).

### Usage notes

As a usage example, here we present the findings in characterizing the age-associated proteome using the proteomic data reported in this work. In this usage showcase, we utilized the entire set of the samples that passed our rigorous QC process (see above), which include 265 novel proteomics profiling (**Supplementary Table 2**), and 274 bulk RNA-seq assays of both new and existing sample sets (**Figure 2A**; **Table 1**).

We first adjusted the protein expression for the covariates including PMI, sex, race, cohort (MSBB vs ROSMAP), and batch (**Figure 4F**). We next calculated the correlation between age at death (AOD) and each of the proteins in the proteome. We detected 1,080 proteins that had a significant correlation with AOD (p.adj < 0.05), which includes 866 and 214 proteins that are negatively and positively associated with AOD, respectively (**Supplementary Table 3**). For example, the ARF4 protein has the highest correlation with AOD (r = −0.44, p.adj = 9.7e-10, Fig. 6A). Previous studies have showed the increased expression of the TMEM106B protein with ageing in human brains^55,56^. This study also revealed that the TMEM106B protein is positively associated with AOD (r = 0.32, p.adj = 1.5e-5, Fig. 6B).

**Figure 6.**
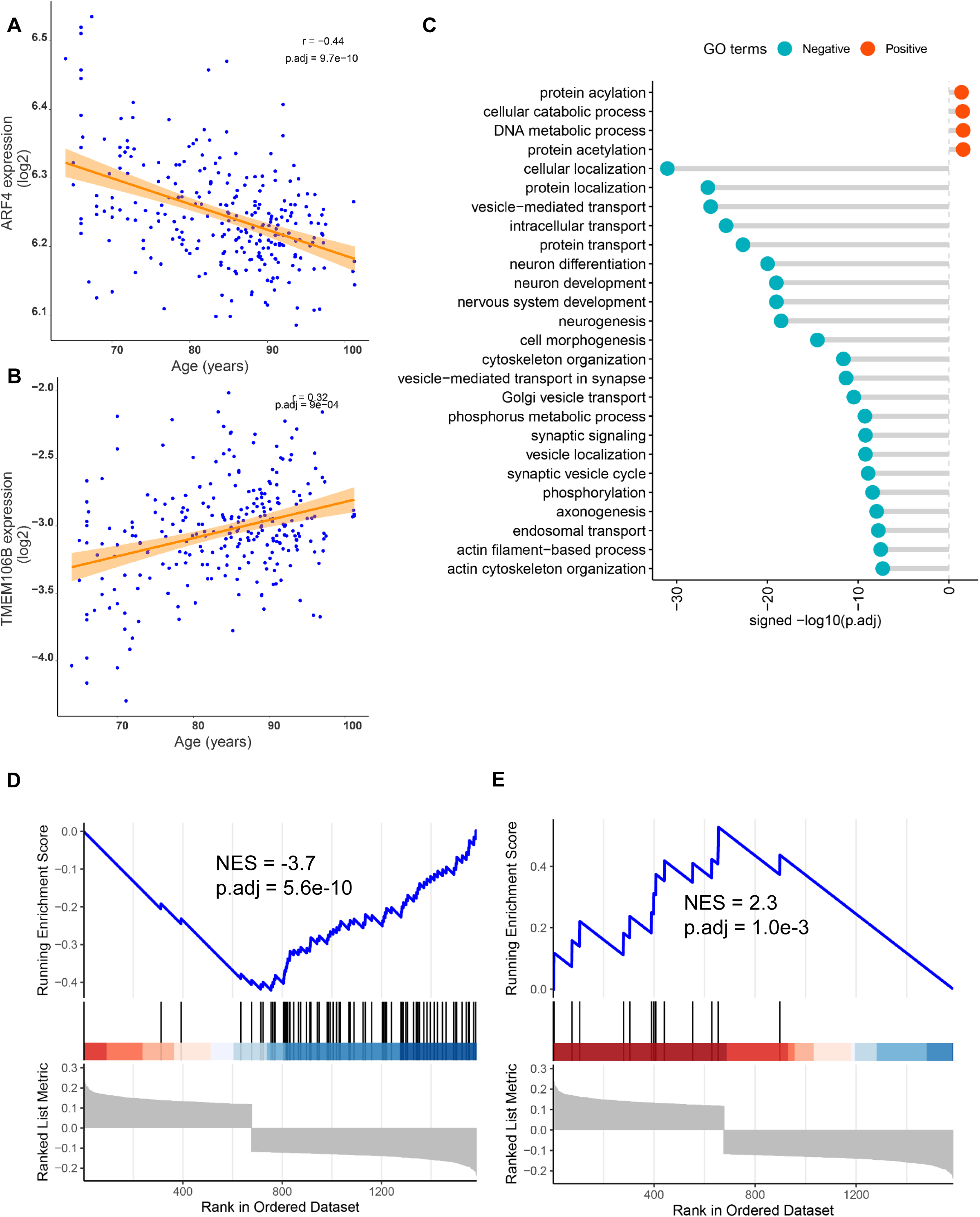
Characterization of age-associated proteome. A-B, scatterplots to show the expression of the ARF4 (A) and TMEM106B (B) protein over age at death. C, dot chart showing enrichment of the age-associated proteins for gene ontology (GO) terms. Negative and Positive represent for Age (at death)-associated proteins negatively and positively, respectively. y-axis, various GO terms. X-axis, -log10(p.adj) for enrichment for each GO term. B, venn diagram to show the overlap between age-associated proteins and genes. Protein(-) and Protein(+) stand for negative and positive age-associated proteins, respectively. Gene(-) and Gene(+) stand for negative and positive age-associated proteins, respectively. FE (fold enrichment) and p.adj stand for fold enrichment and adjusted p value for the overlap, respectively. D-E, the GSEA analysis of the enrichment of the AOD-associated genes for AOD-negatively-and positively-associated protein signatures, respectively. NES, normalized enrichment score. p.adj, adjusted p value.

To find out their biological and functional significance for the AOD-associated protein signatures, we investigated their enrichment for the gene ontology (GO) terms (**Supplementary Table 4**). As shown in **Figure 6C**, the negatively AOD-associated signature are enriched for many biological processes in neuronal and synaptic functions such as neuron development and differentiation, neurogenesis, and synaptic signaling, which is consistent with the observation that this signature is enriched for the neuron-specific gene signature (fold enrichment (FE) = 1.6, and p.adj = 0.008). Additionally, the negatively AOD-associated signatures are enriched for biological processes functioning in cellular transport and cytoskeleton (**Figure 6C**). In contrast, only a few GO terms, for example, cellular catabolic, and DNA metabolic process, were significantly enriched for the positively AOD-associated signature (**Figure 6C**). Our results are in line with the report that GO terms involving synaptic- and cytoskeleton-processes might be an avenue to render the resilience to high AD pathology^3^.

We further tested whether the AOD-associated protein signature can be replicated in the bulk RNA-seq modality. To this end, we performed correlation analysis between AOD and each gene in the transcriptome (see Methods). However, at p.adj < 0.05, only one gene is significantly associated with AOD (**Supplementary Table 5**). Therefore, we chose to use the gene set enrichment analysis (GSEA) to assess the concordance between AOD-associated protein and gene signatures (see Methods). We found that the AOD-associated gene signature (nominal p < 0.05) is negatively and positively enriched in the AOD-negatively- and positively-associated protein signatures, respectively (**Figure 6D-E**), thus, validating the concordance between the AOD-associated protein and gene signatures.

## Availability of data and software code

The MSBB and ROSMAP Cohort AD Resilience Study data are available via the AD Knowledge Portal (https://doi.org/10.7303/9618503). The AD Knowledge Portal is a platform for accessing data, analyses, and tools generated by the Accelerating Medicines Partnership (AMP-AD) Target Discovery Program and other National Institute on Aging (NIA)-supported programs to enable open-science practices and accelerate translational learning. The data, analyses and tools are shared early in the research cycle without a publication embargo on secondary use. Data is available for general research use according to the following requirements for data access and data attribution (https://adknowledgeportal.synapse.org/Data#20Access). All of the codes are available upon request.

## Authors’ contributions

Conceptualization, B.Z.; Investigation, E.W., M.W., L.G., B.X., Y.F., X.W., Z.W., Y.L., X.P., H.L., L.H.; T.Z., C.M.; C.G.; M.E.E.; D.A. B., V.H.; J.P. and B.Z.; Resources, V.H., D.B., J.P., and B.Z.; Writing – Original Draft, E.W., M.W., and B.Z.; Writing – Review & Editing, All Authors; Supervision, J.P. and B.Z.; Funding Acquisition, M.E., V.H., J.P. and B.Z..

## Acknowledgments

We thank Pavel Katsel for his contribution in the collection of brain sample tissues.

